# Deep learning achieves super-resolution in fluorescence microscopy

**DOI:** 10.1101/309641

**Authors:** Hongda Wang, Yair Rivenson, Yiyin Jin, Zhensong Wei, Ronald Gao, Harun Günaydin, Laurent A. Bentolila, Aydogan Ozcan

## Abstract

We present a deep learning-based method for achieving super-resolution in fluorescence microscopy. This data-driven approach does not require any numerical models of the imaging process or the estimation of a point spread function, and is solely based on training a generative adversarial network, which statistically learns to transform low resolution input images into super-resolved ones. Using this method, we super-resolve wide-field images acquired with low numerical aperture objective lenses, matching the resolution that is acquired using high numerical aperture objectives. We also demonstrate that diffraction-limited confocal microscopy images can be transformed by the same framework into super-resolved fluorescence images, matching the image resolution acquired with a stimulated emission depletion (STED) microscope. The deep network rapidly outputs these super-resolution images, without any iterations or parameter search, and even works for types of samples that it was not trained for.

---

Computational super-resolution microscopy techniques in general make use of *a priori* knowledge about the sample and/or the image formation process to enhance the resolution of an acquired image. At the heart of the existing super-resolution methods^1–3^, numerical models are utilized to simulate the imaging process, including, for example, an estimation of the point spread function (PSF) of the imaging system, its spatial sampling rate and/or sensor-specific noise patterns. Fluorescence imaging process is in general more challenging to model and take into account e.g., spatially-varying optical aberrations, the chemical environment of the labeled sample and the optical properties of the specific mounting media and the fluorophores that are used^4–7^. This image modeling related challenge, in turn, leads to formulation of forward models with different simplifying assumptions. In general, more accurate models yield higher quality results, often with a trade-off of exhaustive parameter search and computational cost.

Here we present a deep learning-based framework to achieve super-resolution in fluorescence microscopy without the need for making any assumptions on or precise modeling of the image formation process. Instead, we train a deep neural network using a Generative Adversarial Network (GAN)^8^ model to transform an acquired low-resolution image into a high-resolution one. Once the deep network is trained (see the **Methods** section), it remains fixed and can be used to rapidly output batches of high resolution images, in e.g., 0.4 sec for an image size of 1024×1024 pixels using a single Graphics Processing Unit (GPU). The network inference is noniterative and does not require a manual parameter search to optimize its algorithmic performance. The deep network can also be generalized to different types of samples that were not part of the training process.

We demonstrate the success of this deep learning-based approach by super-resolving the raw images captured by a widefield fluorescence microscope and a confocal microscope. In the widefield imaging case, we transform the images acquired using a 10×/0.4NA objective lens into super-resolved images that match the images of the same samples acquired with a 20×/0.75NA objective lens. In the second case, we transform diffraction-limited confocal microscopy^9^ images to match the resolution of the images that were acquired using a STED microscope^10,11^, showing a PSF width that is improved from ~290 nm down to ~110 nm (i.e., 2.6× improvement). This deep learning-based fluorescence super-resolution framework improves both the field-of-view and throughput of modern fluorescence microscopy tools and can be used to transform low-resolution and wide-field images acquired using various imaging hardware into higher resolution ones.

Recently, a number of studies have used deep learning-based approaches to advance optical microscopy techniques, including bright-field microscopy^12,13^, holographic phase microscopy^14–16^ and fluorescence microscopy.^17–20^ Some of these earlier results on fluorescence microscopy have focused on faster image acquisition or inference for single molecule localization microscopy^17–19^, or resolution enhancement by learning a sample specific imaging process through simulations^20^. Unlike these contributions, our presented technique makes no prior assumptions regarding the imaging process, such as an approximate model of the point spread function^17–20^, and does not depend on an additional computational technique to generate the desired target images, using e.g., PSF-fitting to a sparse set of samples.^17–19^ Rather than localizing specific filamentous structures of a sample, here we demonstrate the generalization of our approach by super-resolving various sub-cellular structures, such as nuclei, microtubules, F-actin and mitochondria. We further demonstrate that the presented framework can be generalized to multiple microscopic imaging modalities, including cross-modality image transformations (e.g., confocal to STED) as we report in the **Results** section.

## RESULTS

### Super-resolution of fluorescently-labeled intracellular structures

We initially demonstrated the super-resolution capability of the presented approach by imaging bovine pulmonary artery endothelial cell (BPAEC) structures; the raw images, used as input to the network, were acquired using a 10×/0.4NA objective lens and the results of the network were compared against the ground truth images, which were captured using a 20×/0.75NA objective lens. An example of the network input image is shown in Fig 1(a), where the field-of-view (FOV) of the 10× and 20× objectives are also labeled. Figs. 1(b, e) show some zoomed-in regions-of-interest (ROI) revealing further details of a cell’s F-actin and microtubules. A pretrained deep neural network is applied to each color channel of these input images (10×/0.4NA), outputting the resolution-enhanced images shown in Figs. 1(c, f), where various features of F-actin, microtubules, and nuclei are clearly resolved at the network output image, providing a very good agreement to the ground truth images (20×/0.75NA) shown in Figs. 1(d, g). Note that all the network output images shown in this work were blindly generated by the deep network, i.e., the input images were not previously seen by the network.

**Figure 1.**
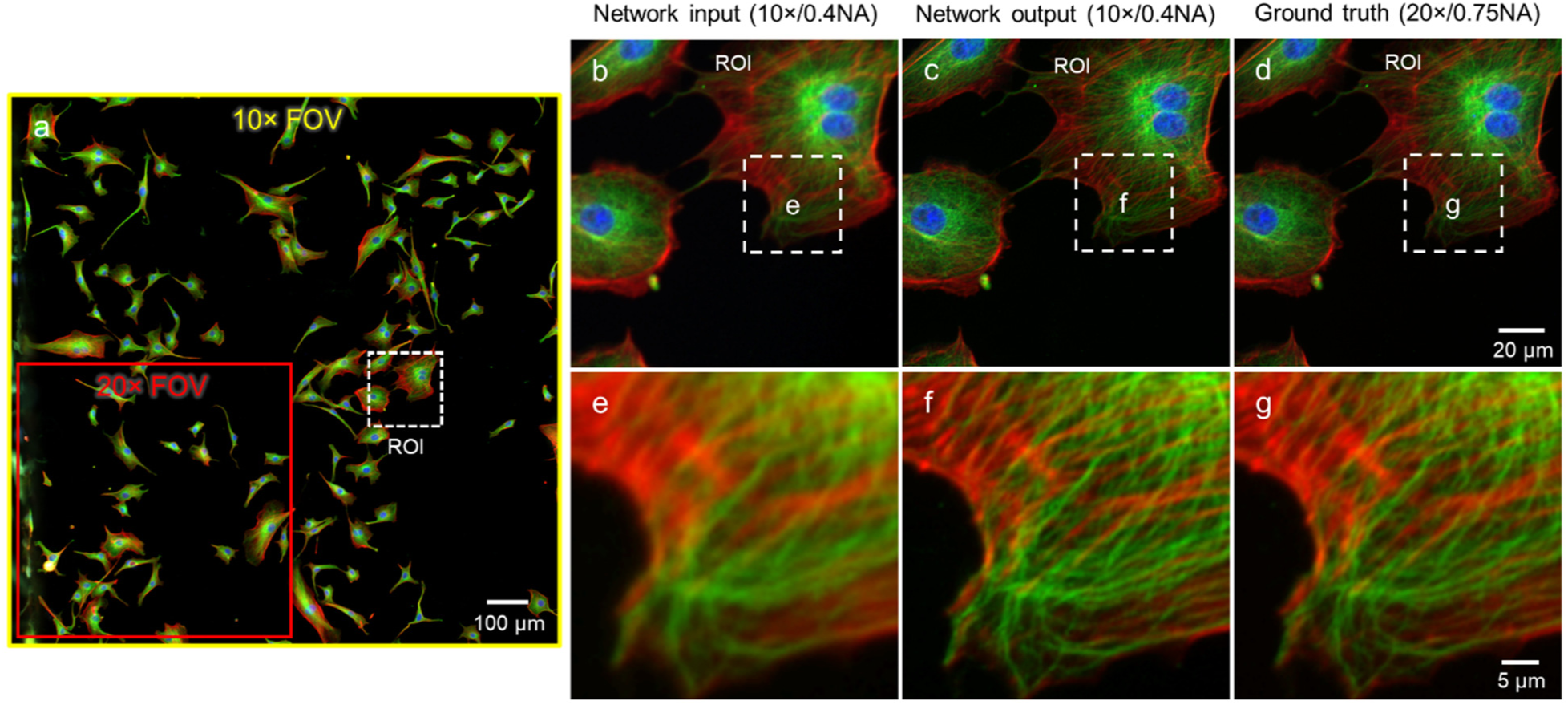
Deep learning-based super-resolved images of bovine pulmonary artery endothelial cells (BPAEC). (a) Network input image acquired with a 10×/0.4NA objective lens. A small ROI is zoomed-in and shown in (b) network input, (c) network output, and (d) ground truth (20×/0.75NA). (e-g) Further zoom-in on a cell’s F-actin and microtubules, corresponding to each image in (d-f).

Next, we compared the results of deep learning-based super-resolution against a widely-used image deconvolution method, i.e., the Lucy-Richardson deconvolution.^21,22^ For this, we used an estimated model of the PSF of the imaging system, which is required by the Lucy-Richardson deconvolution algorithm to approximate the forward model of the image blur. Following its parameter optimization (see the **Methods** Section), the Lucy-Richardson deconvolution algorithm, as expected, demonstrated resolution improvements compared to the input images, as shown in Fig. 2(a-3), (b-3), and (c-3); however compared to deep learning results (Fig. 2(a-2), (b-2), and (c-2)), the improvements observed with Lucy-Richardson deconvolution are modest, despite the fact that it used parameter search/optimization and *a priori* knowledge on the PSF of the imaging system. We also noticed that the deep network output image shows sharper details compared to the ground truth image, especially for the F-actin structures (e.g., Fig. 2(c)). This result is in-line with the fact that all the images were captured by finding the autofocusing plane within the sample using the FITC channel (see e.g., Fig. 2(b-4)), and therefore the Texas-Red channel can remain slightly out-of-focus due to the thickness of the cells. This means the shallow depth-of-field (DOF) of a 20×/0.75NA objective lens (~1.4 μm) might have caused some blurring in the F-actin structures (Fig. 2(c-4)). This out-of-focus imaging of different color channels is not impacting the network output image as much since the input image to the network was captured with a much larger DOF (~5.1 μm), using a 10×/0.4NA objectives lens. Therefore, in addition to an increased FOV resulting from a low NA input image, the network output image is also benefiting from an increased DOF, helping to reveal some finer features that might be out-of-focus in different color channels using a high NA objective lens.

**Figure 2.**
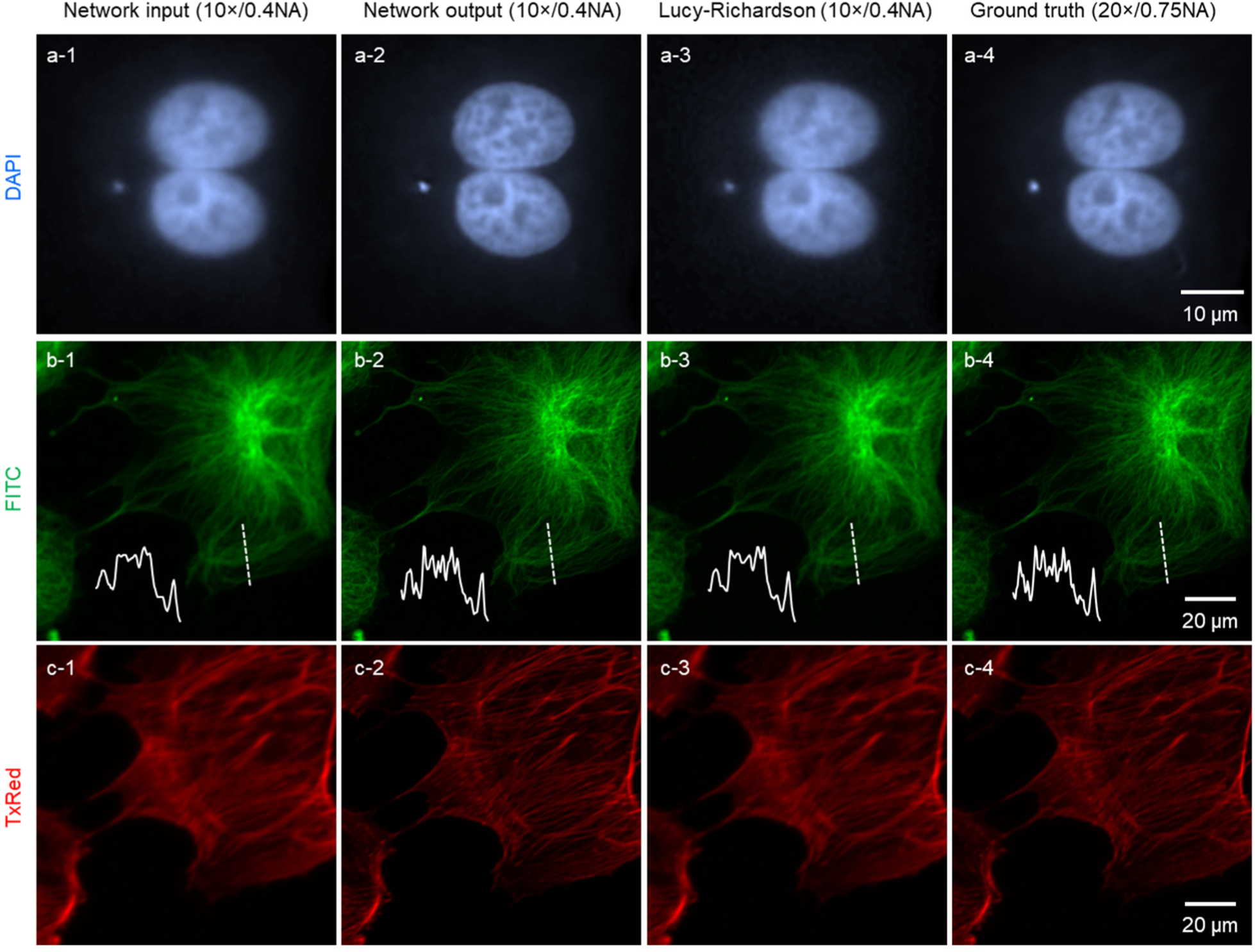
Comparison of deep learning results against Lucy-Richardson image deconvolution.

Next, we tested the generalization of our pre-trained network model in improving image resolution on new types of samples that were not present in the training phase. Figs. 3(a-c) demonstrate the resolution enhancement of mitochondria labeled with MitoTracker Red CMXRos by using a deep neural network that was trained with only the images of F-actin labeled with Texas Red-X phalloidin. Even though such mitochondrial structures were not part of the network’s training set, the deep network was able to correctly infer these structures in its blind inference. Another example is shown in Figs. 3(d-f): the F-actin structure labeled with Alexa Fluor 488 phalloidin is super-resolved by a neural network that was pre-trained with only the images of microtubules labeled with BODIPY FL. These results highlight that our neural network does not overfit to a specific type of structure or specimen, but learns to generalize the transformation between two different fluorescence imaging conditions, which will be further discussed in the **Discussion** section.

**Figure 3.**
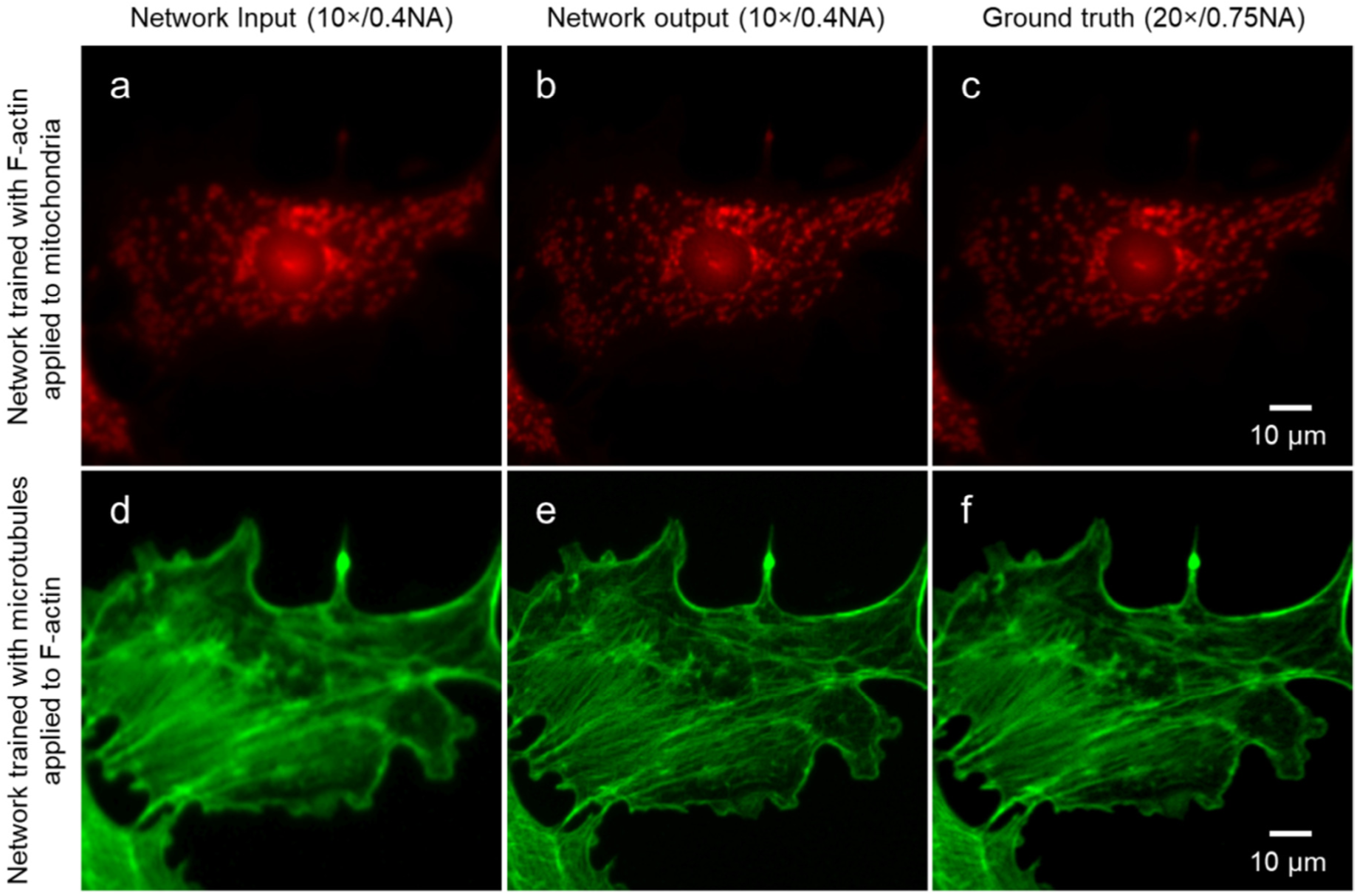
Generalization of a pre-trained neural network model to new types of structures that it was not trained for. (a) Network input, (b) network output, and (c) ground truth images show that the mitochondria inside a BPAEC can be super-resolved by a neural network that was trained with only F-actin images. (d) Network input, (e) network output, and (f) ground truth images show that F-actin inside a BPAEC can be super-resolved by a neural network that was trained with only microtubule images.

### Super-resolution from confocal to STED images

In addition to wide-field fluorescence microscopy, we also applied the presented framework to transform confocal microscopy images into images that match the resolution of STED microscopy; these results are summarized in Figs. 4 and 5, where 20 nm fluorescent beads with 645 nm emission wavelength were imaged on the same platform using both a confocal microscope and a STED microscope (see the **Methods** section). After the training phase, the neural network, as before, blindly takes an input image (confocal) and outputs a super-resolved image that matches the STED image of the same region of interest. Some of the nano-beads in our samples were spaced close to each other, within the classical diffraction limit, i.e., under ~290 nm, as shown in e.g., Fig. 4(d-f), and therefore could not be resolved in the raw confocal microscopy images. The neural network super-resolved these closely-spaced nano-particles, providing a good match to STED images of the same regions of the sample, see Figs. 4(g, h, i) vs. 4(j, k, l).

**Figure 4.**
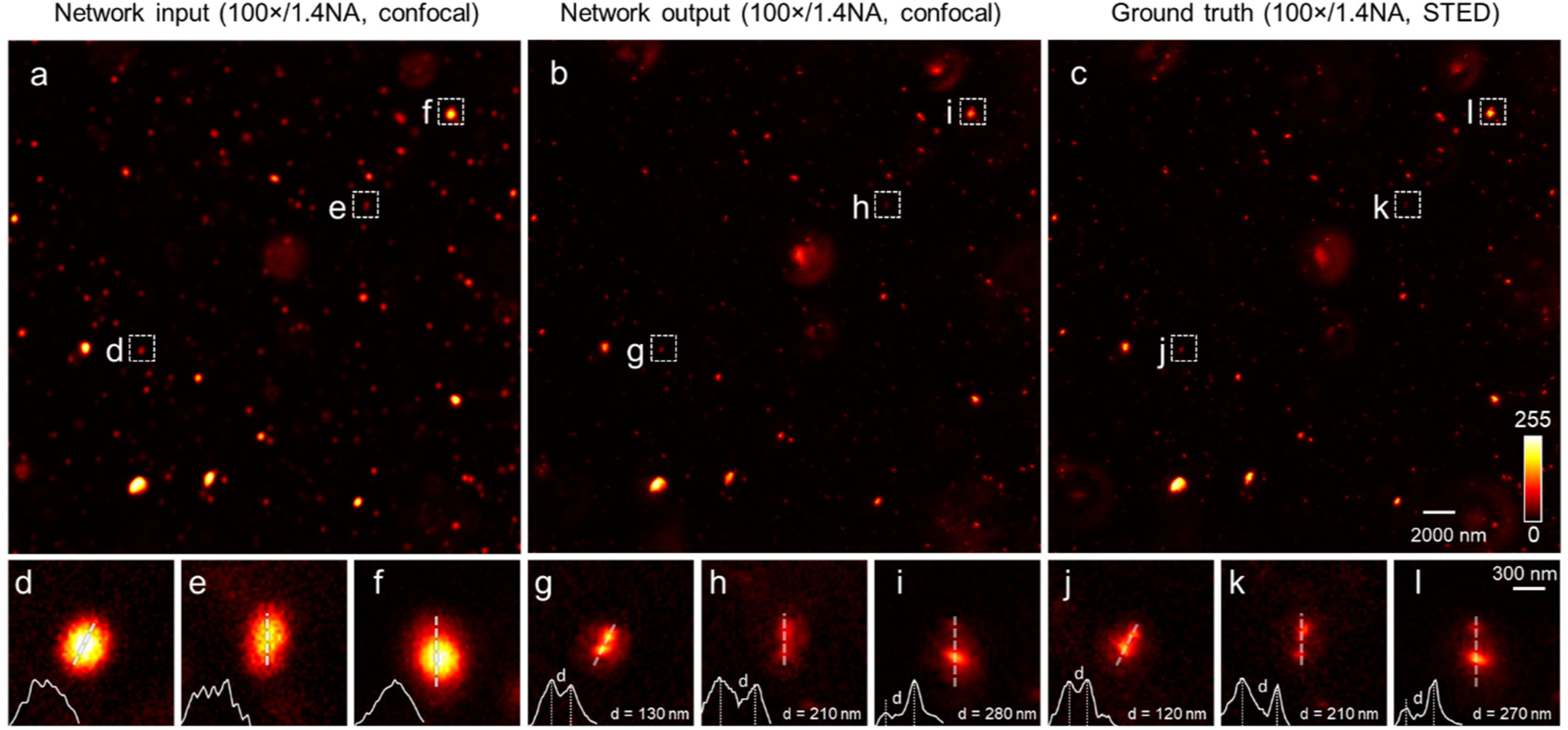
Image resolution improvement beyond the diffraction limit: from confocal microscopy to STED. (a) A diffraction-limited confocal microscope image is used as input to the network and is super-resolved to blindly yield (b) the network output, which is comparable to (c) STED image of the same FOV, used as the ground truth. (d-f) show examples of closely spaced nanobeads that cannot be resolved by confocal microscopy. (h-i) The trained neural network takes (d-f) as input and resolves the individual beads, very well agreeing with (j-l) STED microscopy images. The cross-sectional profiles reported in (d-l) are extracted from the original images. Also see Fig. 5 for further quantification of the performance of the deep network on confocal images, and its comparison to STED.

**Figure 5.**
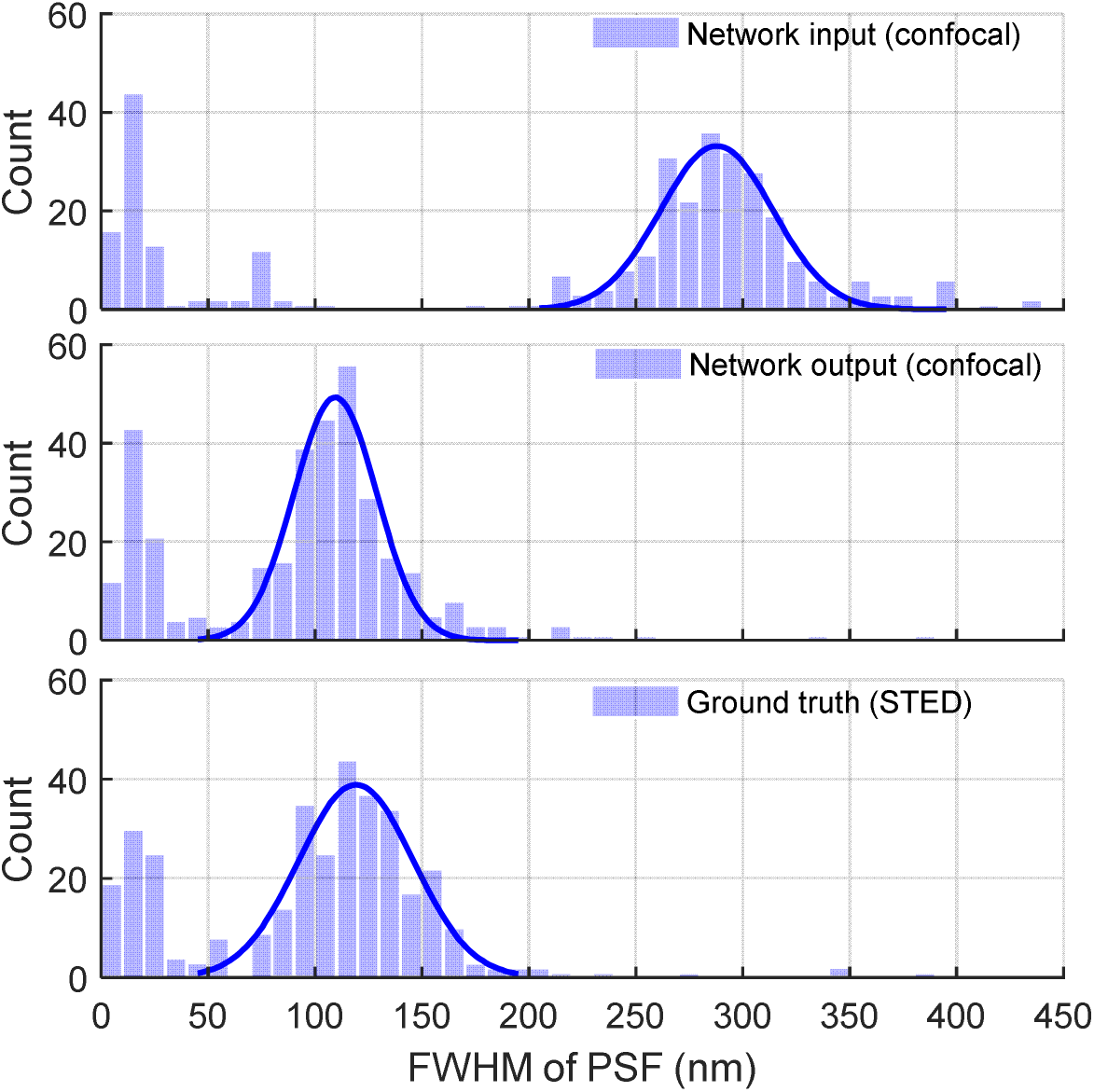
PSF characterization, before and after the network, and its comparison to STED. We extracted more than 400 bright spots from the same locations of the network input (confocal), network output (confocal), and the corresponding ground truth (STED) images. Each one of these spots was fit to a 2D Gaussian function and the corresponding FWHM distributions are shown in each histogram. These results show that the resolution of the network output images is significantly improved from ~290 nm (top row: network input using a confocal microscope) down to ~110 nm (middle row: network output), which provides a very good fit to the ground truth STED images of the same nano-particles, summarized at the bottom row.

To further quantify this resolution improvement achieved by the neural network, we measured the PSFs arising from the images of single/isolated nano-beads across the sample field-of-view^23^; this was repeated for more than 400 individual particles that were tracked in the images of the confocal microscope and STED microscope, as well as the network output image (in response to the confocal image). The results are summarized in Fig. 5, where the full-width half-maximum (FWHM) of the confocal microscope PSF is centered at ~290 nm, roughly corresponding to the lateral resolution of a diffraction limited imaging system at an emission wavelength of 645 nm. As shown in Fig. 5, PSF FWHM distribution of the network output provides a very good match to the PSF results of the STED system, with a mean FWHM of ~110 nm vs. ~120 nm, respectively.

## DISCUSSION

The generalized point spread function of an imaging system, which accounts for the finite aperture of the optical system, as well as its aberrations, noise and optical diffraction, can be considered as a probability density function, *p*(*ζ*,*η*), where *ζ*, *η* denote the spatial coordinates. *p* (*ζ*,*η*) represents the probability of photons emitted from an ideal point source on the sample to arrive at a certain displacement on the detector plane. Therefore, the super-resolution task that the presented deep learning framework has been learning is to transform the input data distribution *X* (*p_LR_* (*ζ*, *η*)) into a high-resolution output, *Y* (*p_HR_* (*ζ*, *η*)), where the former is created by a lower resolution (LR) imaging system and the latter represents a higher resolution (HR) imaging system. The network architecture that we have used for training, i.e., GANs^8^ have been proven to be extremely effective in learning such distribution transformations (*X* → *Y*) *without* any prior information on or modelling of the image formation process or its parameters.^24,25^ Unlike other statistical super-resolution methods, the presented approach is data-driven, and the deep network is trying to find a distribution generated by real microscopic imaging systems that it was trained with. This feature makes the network more resilient to poor image SNR (signal-to-noise ratio) and related challenges, and the presented method is not susceptible to aberrations of the imaging parameters, such as the PSF^5^ and sensor-specific noise patterns, which are required for any standard deconvolution and localization method^26^. A similar resilience to spatial and spectral aberrations of an imaging system has also been demonstrated for bright-field microscopic imaging using a neural network.^12^

The capability of transforming a fluorescence microscopic image into a higher resolution one not only shortens the image acquisition time because of the increased FOV of low NA systems, but also enables new opportunities for imaging objects that are vulnerable to photo-bleaching or photo-toxicity.^27,28^ For example, in the experiments reported in Figs. 4 and 5, the required excitation power for STED microscopy was 10-fold stronger than that of confocal microscopy, as detailed in the **Methods** section. Furthermore, the depletion beam of STED microscopy is typically orders of magnitude higher than its excitation beam, which sets practical challenges for some biomedical imaging applications.^28–30^ Most of these issues become less pronounced when using a confocal microscopy system, which is also quite simpler in its hardware compared to a STED microscope.^31^ Using the presented deep learning-based approach, the diffraction induced resolution gap between a STED image and a confocal microscope image can be closed, achieving super-resolution microscopy using relatively simpler and more cost-effective imaging systems, also reducing photo-toxicity and photo-bleaching.

Another important feature of the deep network-based super-resolution approach is that it can resolve features over an extended DOF because a low NA objective is used to acquire the input image; see e.g., Fig. 2(c) and Fig. 3(e, f) for the F-actin structures. A similar observation was also made for deep learning-enhanced bright-field microscopy images reported earlier.^13^ This extended DOF is also favorable in terms of photo-damage to the sample, by eliminating the need for a fine axial scan within the sample volume, which might reduce the overall light delivered to the sample, while making the imaging process more efficient.

A common concern for computational approaches that enhance image resolution is the potential emergence of spatial artifacts which may degrade the image quality, such as the Gibbs phenomenon in Lucy-Richardson deconvolution.^32^ To explore this, we randomly selected an example in the test image dataset, and quantified the artifacts of the network output image using the NanoJ-Squirrel Plugin^5^; this analysis revealed that the network output image does not generate noticeable super-resolution artifacts and in fact has the same level of spatial mismatch error that the ground truth HR image has with respect to the LR input image of the same sample (see **Supplementary** Fig. S1 and **Supplementary** Note 1). This conclusion is further confirmed by **Supplementary** Fig. S1(d), which overlays the network output image and the ground truth image, revealing no obvious feature mismatch between the two. The same conclusion remained consistent for other test images as well. Since our deep network models are trained within the GAN framework, potential image artifacts and hallucinations of the generative network were continuously being suppressed and accordingly penalized by the discriminative model during the training phase, which helped the final generative network to be robust and realistic in its superresolution inference. Moreover, in case feature hallucinations are observed in e.g., the images of new types of samples, these can be additionally penalized in the loss function as they are discovered, and the network can be further regularized to avoid such artifacts from repeating.

## METHODS

### Wide-field fluorescence microscopic image acquisition

The fluorescence microscopic images (Figs. 1 and 2) were captured by scanning a microscope slide containing multi-labeled bovine pulmonary artery endothelial cells (BPAEC) (FluoCells Prepared Slide #2, Thermo Fisher Scientific) on a standard inverted microscope which is equipped with a motorized stage (IX83, Olympus Life Science). The low-resolution (LR) and high-resolution (HR) images were acquired using 10×/0.4NA (UPLSAPO10X2, Olympus Life Science) and 20×/0.75NA (UPLSAPO20X, Olympus Life Science) objective lenses, respectively. Three bandpass optical filter sets were used to image the three different labelled cell structures and organelles: Texas Red for F-actin (OSFI3-TXRED-4040C, EX562/40, EM624/40, DM593, Semrock), FITC for microtubules (OSFI3-FITC-2024B, EX485/20, EM522/24, DM506, Semrock), and DAPI for nuclei (OSFI3-DAPI-5060C, EX377/50, EM447/60, DM409, Semrock). The imaging experiments were controlled by MetaMorph microscope automation software (Molecular Devices), which performed translational scanning and auto-focusing at each position of the stage. The auto-focusing was performed on the FITC channel, and the DAPI and Texas Red channels were both exposed at the same plane as FITC. With a 130 W fluorescence light source set to 25% output power (U-HGLGPS, Olympus Life Science), the exposure time for each channel was set to: Texas Red 350 ms (10×) and 150 ms (20×), FITC 800 ms (10×) and 400 ms (20×), DAPI 60 ms (10×) and 50 ms (20×). The images were recorded by a monochrome scientific CMOS camera (ORCA-flash4.0 v2, Hamamatsu Photonics K.K.) and saved as 16-bit grayscale images with regards to each optical filter set. The additional test images (Fig. 3) are captured using the same setup with FluoCells Prepared Slide #1 (Thermo Fisher Scientific), with the filter setting of Texas Red for mitochondria, and FITC for F-actin.

### Confocal and STED image acquisition

The samples for confocal and STED experiments (Figs. 4, 5) were prepared with 20 nm fluorescent nano-beads (FluoSpheres Carboxylate-Modified Microspheres, crimson fluorescent (625/645), 2% solids, Thermo Fisher Scientific) that were diluted 100 times with methanol and sonicated for 3×10 minutes, and then mounted with antifade reagents (ProLong Diamond, Thermo Fisher Scientific) on a standard glass slide, followed by placing a 0.17 mm-thick cover glass (Carl Zeiss Microscopy). The confocal and STED imaging experiments were performed on a laser scanning confocal and STED microscope (TCS SP8, controlled by Leica Application Suite X, Leica Microsystems) with a 100×/1.4NA oil immersion objective lens (HC PL APO 100x/1.4 OIL CS2, Leica Microsystems). The scanning for each FOV was performed by a resonant scanner working at 8000 Hz with 16 times line average and 30 times frame average. The fluorescent nano-beads were excited with a laser beam at 633 nm wavelength. The emission signal was captured with a hybrid photodetector (HyD SMD, Leica Microsystems) with 440 V active gain through a 645~752 nm bandpass filter. The excitation laser power was set to 5% for confocal imaging, and 50% for STED imaging, so that the signal intensities remained similar while keeping the same scanning speed and gain voltage. A depletion beam of 775 nm was also applied when capturing STED images with 100% power. The confocal pinhole was set to 1 Airy unit (e.g., 168.6 μm for 645 nm emission wavelength and 100× magnification) for both the confocal and STED imaging experiments. The scanning step size (i.e., the effective pixel size) was ~30.4 nm to ensure sufficient sampling rate. All the images were exported and saved as 8-bit grayscale images.

### Image pre-processing

For widefield images (Figs. 1, 2, and 3), a low intensity threshold was applied to subtract background noise and auto-fluorescence, as a common practice in fluorescence microscopy. The threshold value was estimated from the mean intensity value of a region without objects, which is ~300 out of 65535 in our 16-bit images. The LR images are then linearly interpolated two times to match the effective pixel size of the HR images. Accurate registration of the corresponding LR and HR training image pairs is of crucial importance since the objective function of our network consists of adversarial loss and pixel-wise loss. We employed a two-step registration workflow to achieve the needed registration with sub-pixel level accuracy (see **Supplementary** Fig. S2). First, the fields-of-view of LR and HR images are digitally stitched in a MATLAB script interfaced with Fiji^33^ Grid/Collection stitching plugin^34^ through MIJ^35^, and matched by fitting their normalized cross-correlation map to a 2D Gaussian function and finding the peak location (see **Supplementary** Note 2). However, due to the optical distortion and color aberration of different objective lenses, the local features might still not be exactly matched. To address this, the globally matched images are fed into a pyramidal elastic registration algorithm to achieve sub-pixel level matching accuracy, which is an iterative version of the registration module in Fiji Plugin NanoJ, with a shrinking block size (see **Supplementary** Fig. S2).^5,12,24,33^ This registration step starts with a block size of 256×256 and stops at a block size of 64×64, while shrinking the block size by 1.2 times every 5 iterations with a shift tolerance of 0.2 pixels. Due to the slightly different placement and the distortion of the optical filter sets, we performed the pyramidal elastic registration for each fluorescence channel independently. At the last step, the precisely registered images were cropped 10 pixels on each side to avoid registration artifacts, and converted to single-precision floating data type and scaled to a dynamic range of 0~255. This scaling step is not mandatory but creates convenience for fine tuning of hyperparameters when working with images from different microscopes/sources.

For confocal and STED images (Figs. 4, 5) which were scanned in sequence on the same platform, only a drift correction step was required, which was calculated from the 2D Gaussian fit of the cross-correlation map. The drift was found to be ~10 nm for each scanning FOV between the confocal and STED images. We did not perform thresholding to this dataset for the network training. However, after the test images were enhanced by the network, we subtracted a constant value (calculated by taking the mean value of an empty FOV) from the confocal (network input), the super-resolved (network output), and the STED (ground truth) images, respectively, for better visualization and comparison of the images. The total number of images used for training, validation and blind testing of each network are summarized in Supplementary Table 1.

### Generative adversarial network structure and training

In this work, our deep neural network was trained following the generative adversarial network (GAN) framework^8^, which has two sub-networks being trained simultaneously, a generative model which enhances the input LR image, and a discriminative model which returns an adversarial loss to the resolution-enhanced image, as illustrated in Fig. 6. We designed our objective function as the combination of the adversarial loss with two regularization terms: the mean square error (MSE), and the structural similarity (SSIM) index^36^. Specifically, we aim to minimize:

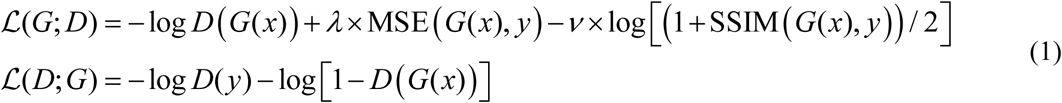

where *x* is the LR input, *G* (*x*) is the generative model output, *D* (•) is the discriminative model prediction of an image (network output or ground truth image), and *y* is the HR image used as ground truth. The structural similarity index is defined as:

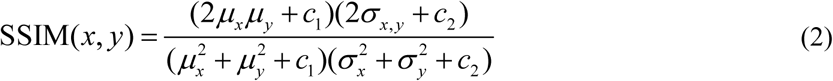

where *μ_x_*, *μ_y_* are the averages of 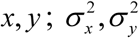 are the variances of *x*, *y*; *σ_x,y_* is the covariance of *x* and *y*; and *c*_l_, *c*_2_ are the variables used to stabilize the division with a small denominator. An SSIM value of 1.0 refers to identical images. When training with the wide-field fluorescence images, the regularization constants *λ* and *v* were set to accommodate the MSE loss and the SSIM loss to be ~1% of the combined generative model loss *ℒ*(*G*;*D*). When training with the confocal-STED image datasets, we kept *λ* the same and set *v* to 0. While the adversarial loss guides the generative model to map the LR images into HR, the two regularization terms assure that the generator output image is established on the input image with matched intensity profile and structural features. These two regularization terms also help us stabilize the training schedule and smoothen out the spikes on the training loss curve before it reaches equilibrium. For the subnetwork models, we employed a similar network structure as described in Ref. [24].

**Figure 6.**
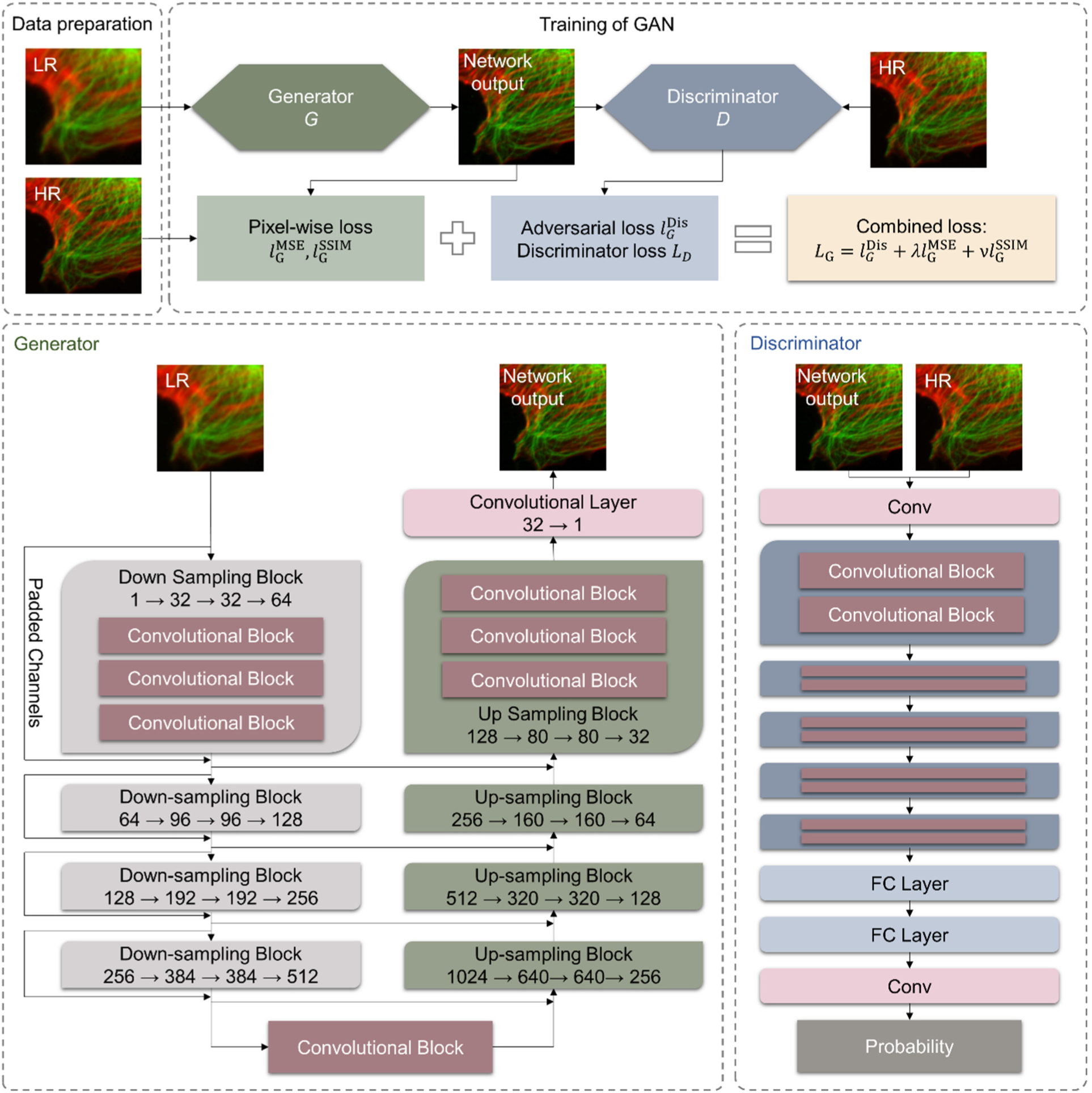
The training process and the architecture of the generative adversarial network that we used for super-resolution.

### Generative Model

U-net is a CNN (convolutional neural network) architecture, which was first proposed for medical image segmentation, yielding high performance with very few training datasets.^37^ The same network architecture has also been successfully applied in recent image reconstruction and virtual staining applications^16,24^. The structure of the generative network used in this work is illustrated in Fig. 6, which consists of four down-sampling blocks and four up-sampling blocks. Each down-sampling block consists of three residual convolutional blocks, within which it performs:

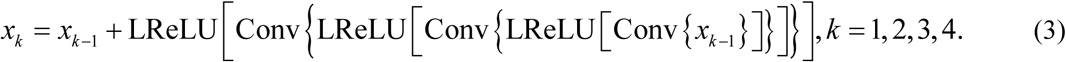

where *x_k_* represents the output of the *k-th* down-sampling block, and *x*_0_ is the LR input image. Conv{ } is the convolution operation, LReLU[ ] is the leaky rectified linear unit activation function with a slope of *α* = 0.1, i.e.,

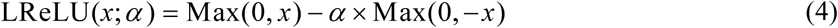

The input of each down-sampling block is zero-padded and added to the output of the same block. The spatial down-sampling is achieved by an average pooling layer after each downsampling block. A convolutional layer lies at the bottom of this U-shape structure that connects the down-sampling and up-sampling blocks.

Each up-sampling block also consists of three convolutional blocks, within which it performs:

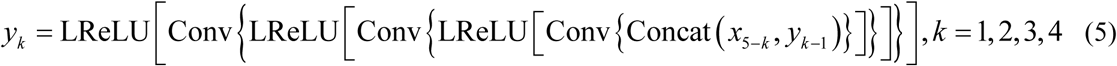

where *y_k_* represents the output of the *k*-th up-sampling block, and *y*_0_ is the input of the first upsampling block. Concat( ) is the concatenation operation of the down-sampling block output and the up-sampling block input on the same level in the U-shape structure. The last layer is another convolutional layer that maps the 32 channels into 1 channel that corresponds to a monochrome grayscale image.

### Discriminative Model

As shown in Fig 6, the structure of the discriminative model begins with a convolutional layer, which is followed by 5 convolutional blocks, each of which performs the following operation:

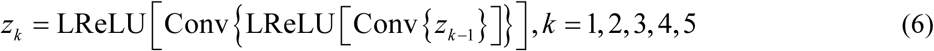

where *z_k_* represents the output of the *k*-th convolutional block, and *z*_0_ is the input of the first convolutional block. The output of the last convolutional block is fed into an average pooling layer whose filter shape is the same as the patch size, i.e., *H* × *W*. This layer is followed by two fully connected layers for dimension reduction. The last layer is a sigmoid activation function whose output is the probability of an input image being ground truth, defined as:

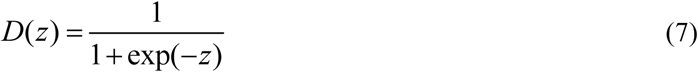

### Network training schedule

During our training the patch size is set to be 64 × 64, with a batch size of 12 on each of the two GPUs. Within each iteration, the generative model and the discriminative model are each updated once while keeping the other unchanged. Both the generative model and the discriminative model were randomly initialized and optimized using the adaptive moment estimation (Adam) optimizer^38^ with a starting learning rate of 1 × 10^−4^ and 1 × 10^−5^, respectively. This framework was implemented with TensorFlow framework version 1.7.0^39^ and Python version 3.6.4 in Microsoft Windows 10 operating system. The training was performed on a consumer grade laptop (EON17-SLX, Origin PC) equipped with dual GeForce GTX1080 graphic cards (NVDIA) and a Core i7-8700K CPU @ 3.7GHz (Intel). The final model for widefield images were selected with the smallest validation loss at around ~50,000^th^ iteration, which took ~10 hours to train. The final model for confocal-STED transformation is selected with the smallest validation loss at around ~500,000^th^ iteration, which took ~90 hours to train.

### Implementation of Lucy-Richardson deconvolution

To make a fair comparison, the lower resolution images were up-sampled 2 times by bilinear interpolation before being deconvolved. We used the Born and Wolf PSF model^40,41^, with parameters set to match our experimental setup, i.e., NA = 0.4, immersion refractive index = 1.0, pixel size = 325 nm. The PSF is generated by an Fiji PSF Generator Plugin^33,42^. We performed an exhaustive parameters search by running the Lucy-Richardson algorithm with 1~100 iterations and damping threshold 0%~10%. The results were visually assessed, with the best one obtained at 10 iterations and 0.1% damping threshold (Fig. 2, third column). The deconvolution for Texas Red, FITC, and DAPI channels were performed separately, assuming central emission wavelengths to be 630 nm, 532nm, and 450 nm, respectively.

### Characterization of the lateral resolution by PSF fitting

The resolution differences among the network input (confocal), the network output (confocal), and the ground truth (STED) images were characterized by fitting their PSFs to a 2D Gaussian profile, as shown in Fig. 5. To do so, more than 400 independent bright spots were selected from the ground truth STED images and cropped out with the surrounding 19×19-pixel regions, i.e., ~577×577 nm^2^. The same locations were also projected to the network input and output images, followed by cropping of the same image regions as in the ground truth STED images. Each cropped region was then fitted to a 2D Gaussian profile. The FWHM values of all these 2D profiles were plotted as histograms, shown in Fig. 5. For each category of images, the histogram profile within the main peak region is fitted to a 1D Gaussian function (Fig. 5).

## ACKNOWLEDGEMENTS

The Ozcan Research Group at UCLA acknowledges the support of NSF Engineering Research Center (ERC, PATHS-UP), the Army Research Office (ARO; W911NF-13-1-0419 and W911NF-13-1-0197), the ARO Life Sciences Division, the National Science Foundation (NSF) CBET Division Biophotonics Program, the NSF Emerging Frontiers in Research and Innovation (EFRI) Award, the NSF INSPIRE Award, NSF Partnerships for Innovation: Building Innovation Capacity (PFI:BIC) Program, the National Institutes of Health (NIH, R21EB023115), the Howard Hughes Medical Institute (HHMI), Vodafone Americas Foundation, the Mary Kay Foundation, and Steven & Alexandra Cohen Foundation. Yair Rivenson is partially supported by the European Union’s Horizon 2020 research and innovation programme under the Marie Skłodowska-Curie grant agreement No H2020-MSCA-IF-2014-659595 (MCMQCT). Confocal and STED laser scanning microscopy was performed at the California NanoSystems Institute (CNSI) Advanced Light Microscopy/Spectroscopy Shared Resource Facility at UCLA.

## REFERENCES

1. Henriques, R. et al. QuickPALM: 3D real-time photoactivation nanoscopy image processing in ImageJ. Nat. Methods 7, 339–340 (2010).

2. Small, A. & Stahlheber, S. Fluorophore localization algorithms for super-resolution microscopy. Nat. Methods 11, 267–279 (2014).

3. Abraham, A. V., Ram, S., Chao, J., Ward, E. S. & Ober, R. J. Quantitative study of single molecule location estimation techniques. Opt. Express 17, 23352–23373 (2009).

4. Dempsey, G. T., Vaughan, J. C., Chen, K. H., Bates, M. & Zhuang, X. Evaluation of fluorophores for optimal performance in localization-based super-resolution imaging. Nat. Methods 8, 1027–1036 (2011).

5. Culley, S. et al. Quantitative mapping and minimization of super-resolution optical imaging artifacts. Nat. Methods 15, 263–266 (2018).

6. Sage, D. et al. Quantitative evaluation of software packages for single-molecule localization microscopy. Nat. Methods 12, 717–724 (2015).

7. Almada, P., Culley, S. & Henriques, R. PALM and STORM: Into large fields and high-throughput microscopy with sCMOS detectors. Methods 88, 109–121 (2015).

8. Goodfellow, I. J. et al. Generative Adversarial Networks. ArXiv14062661 Cs Stat (2014).

9. Wilson, T. & Masters, B. R. Confocal microscopy. Appl. Opt. 33, 565–566 (1994).

10. Betzig, E. et al. Imaging Intracellular Fluorescent Proteins at Nanometer Resolution. Science 313, 1642–1645 (2006).

11. Hell, S. W. & Wichmann, J. Breaking the diffraction resolution limit by stimulated emission: stimulated-emission-depletion fluorescence microscopy. Opt. Lett. 19, 780–782 (1994).

12. Rivenson, Y. et al. Deep learning enhanced mobile-phone microscopy. ArXiv171204139 Phys. (2017).

13. Rivenson, Y. et al. Deep learning microscopy. Optica 4, 1437–1443 (2017).

14. Rivenson, Y., Zhang, Y., Günaydın, H., Teng, D. & Ozcan, A. Phase recovery and holographic image reconstruction using deep learning in neural networks. Light Sci. Appl. 7, 17141 (2018).

15. Sinha, A., Lee, J., Li, S. & Barbastathis, G. Lensless computational imaging through deep learning. Optica 4, 1117–1125 (2017).

16. Wu, Y. et al. Extended depth-of-field in holographic image reconstruction using deep learning based auto-focusing and phase-recovery. (2018).

17. Boyd, N., Jonas, E., Babcock, H. P. & Recht, B. DeepLoco: Fast 3D Localization Microscopy Using Neural Networks. bioRxiv 267096 (2018).doi:10.1101/267096

18. Ouyang, W., Aristov, A., Lelek, M., Hao, X. & Zimmer, C. Deep learning massively accelerates super-resolution localization microscopy. Nat. Biotechnol. (2018). doi:11038/nbt.4106

19. Nehme, E., Weiss, L. E., Michaeli, T. & Shechtman, Y. Deep-STORM: super-resolution single-molecule microscopy by deep learning. Optica 5, 458–464 (2018).

20. Weigert, M. et al. Content-Aware Image Restoration: Pushing the Limits of Fluorescence Microscopy. bioRxiv 236463 (2018).doi:11101/236463

21. Richardson, W. H. Bayesian-Based Iterative Method of Image Restoration. J. Opt. Soc. Am. 1917–1983 62, 55 (1972).

22. Lucy, L. B. An iterative technique for the rectification of observed distributions.Astron. J. 79, 745 (1974).

23. Farahani, J. N., Schibler, M. J. & Bentolila, L. A. Stimulated emission depletion (STED) microscopy: from theory to practice. Microsc. Sci. Technol. Appl. Educ. 2, 1539–1547 (2010).

24. Rivenson, Y. et al. Deep learning-based virtual histology staining using auto-fluorescence of label-free tissue. ArXiv180311293 Phys. (2018).

25. Sønderby, C. K., Caballero, J., Theis, L., Shi, W. & Huszár, F. Amortised MAP Inference for Image Super-resolution. ArXiv161004490 Cs Stat (2016).

26. Li, Y. et al. Real-time 3D single-molecule localization using experimental point spread functions. Nat. Methods (2018). doi:11038/nmeth.4661

27. Dyba, M. & Hell, S. W. Photostability of a fluorescent marker under pulsed excited-state depletion through stimulated emission. Appl. Opt. 42, 5123–5129 (2003).

28. Wäldchen, S., Lehmann, J., Klein, T., Linde, S. van de & Sauer, M. Light-induced cell damage in live-cell super-resolution microscopy. Sci. Rep. 5, 15348 (2015).

29. Hein, B., Willig, K. I. & Hell, S. W. Stimulated emission depletion (STED) nanoscopy of a fluorescent protein-labeled organelle inside a living cell. Proc. Natl. Acad. Sci. 105, 14271–14276 (2008).

30. Hein, B. et al. Stimulated Emission Depletion Nanoscopy of Living Cells Using SNAP-Tag Fusion Proteins. Biophys.J. 98, 158–163 (2010).

31. Brelje, T. C., Wessendorf, M. W. & Sorenson, R. L. Chapter 4 Multicolor Laser Scanning Confocal Immunofluorescence Microscopy: Practical Application and Limitations. in Methods in Cell Biology (ed. Matsumoto, B.) 38, 97–181 (Academic Press, 1993).

32. Liu, R. & Jia, J. Reducing boundary artifacts in image deconvolution. in 2008 15th IEEE International Conference on Image Processing 505–508 (2008).doi:11109/ICIP.2008.4711802

33. Schindelin, J. et al. Fiji: an open-source platform for biological-image analysis. Nat. Methods 9, 676–682 (2012).

34. Preibisch, S., Saalfeld, S. & Tomancak, P. Globally optimal stitching of tiled 3D microscopic image acquisitions. Bioinformatics 25, 1463–1465 (2009).

35. Sage, D., Prodanov, D., Tinevez, J.-Y. & Schindelin, J. MIJ: Making interoperability between ImageJ and Matlab possible. in ImageJ User & Developer Conference (2012).

36. Wang, Z., Bovik, A. C., Sheikh, H. R. & Simoncelli, E. P. Image Quality Assessment: From Error Visibility to Structural Similarity. IEEE Trans. Image Process. 13, 600–612 (2004).

37. Ronneberger, O., Fischer, P. & Brox, T. U-Net: Convolutional Networks for Biomedical Image Segmentation. ArXiv150504597 Cs (2015).

38. Kingma, D. P. & Ba, J. Adam: A Method for Stochastic Optimization. ArXiv14126980 Cs(2014).

39. Abadi, M. et al. TensorFlow: A system for large-scale machine learning. ArXiv160508695 Cs (2016).

40. Aguet, F., Ville, D. V. D. & Unser, M. Model-Based 2.5-D Deconvolution for Extended Depth of Field in Brightfield Microscopy. IEEE Trans. Image Process. 17, 1144–1153 (2008).

41. Born, M., Wolf, E. & Bhatia, A. B. Principles of Optics: Electromagnetic Theory of Propagation, Interference and Diffraction of Light. (Cambridge University Press, 1999).

42. Kirshner, H., Aguet, F., Sage, D. & Unser, M. 3-D PSF fitting for fluorescence microscopy: implementation and localization application. J. Microsc. 249, 13–25 (2013).

